# Rapid long-range synaptic remodeling in hyperacute ischemic stroke

**DOI:** 10.1101/2024.12.12.628225

**Authors:** Huanhuan Chen, Ye Wei, Luminiţa Ruje, Qi Wan, Mikhail Spivakov, Oleg O. Glebov

## Abstract

Physiological mechanisms of the key hyperacute (0-24 hours) stage of stroke remain incompletely understood, hampering development of new stroke therapies. Synaptic plasticity has been strongly implicated in early stages of neurodegenerative and neurodevelopmental disorders. Here, we describe rapid region-specific patterns of synaptic remodeling in stroke, arising within 4 hours of onset. While the ischemic core exhibits profound structural and functional synaptic decline, synapses in the mildly ischemic penumbra largely retain their structure, decreasing synaptic function. Conversely, synapses in the contralateral cortex across the brain show widespread structural and functional increase. Mechanistically, hyperacute stroke triggers synaptic recruitment of NMDA receptors in the ischemic core and penumbra, while NMDA receptor blockade exacerbates structural synaptic decline in the penumbra and abolishes contralateral synaptic enhancement. Proteomic analysis confirms cross-brain synaptic strengthening and reveals emergence of metabolic rearrangement in the penumbra, while RNAseq indicates broad downregulation of synaptic gene expression in the penumbra. These findings identify brain-wide homeostatic synaptic rebalancing as a potential mechanism for rapid-response functional compensation in early stroke, highlighting the extent of brain resilience to acute perturbation.

**Highlights:**

- Stroke induces rapid long-range synaptic enhancement in the contralateral hemisphere
- NMDAR signaling orchestrates short and long-range synaptic remodeling in stroke
- Proteomics and RNAseq reveal details of region-specific synaptic remodeling

**eTOC blurb:** What happens to synaptic communication across the brain during the first hours of stroke is unclear. We find that while synapses at the center of the stroke quickly fall apart, those in the surrounding area are protected and those in the opposite hemisphere are rapidly strengthened by the activity of NMDA-type glutamate receptors.

## Introduction

As the second and third leading cause of mortality and disability respectively, stroke represents a significant burden on public health, with rising global incidence.^1–3^ The initial hyperacute (0-24 hours)^4^ stage of stroke is of particular clinical relevance, as urgent therapeutic intervention to mitigate brain damage is key for preservation of functionality, as reflected in the denomination of this time period as “the golden hour” of stroke.^5–7^ Nevertheless, current therapy options for hyperacute stroke remain limited, focusing exclusively on restoration of the blood flow, while clinical criteria exclude a significant proportion of patients from treatment.^7–9^ Better understanding of molecular, cellular and physiological events during the “golden hour” is therefore sorely needed for future development of new stroke therapeutics.

A major factor underlying central nervous system (CNS) pathology is dysregulation of synaptic function. Synaptic plasticity has been extensively implicated in early etiology of neurodevelopmental and neurodegenerative disorders, becoming a major topic for basic and translational research.^10–15^ In stark contrast, investigation of the synapse in early stroke has seen little progress beyond demonstration of broad synaptic failure in *ex vivo* ischemia models more than 2 decades ago.^16–20^ Crucially, such simplistic models could not provide opportunities for investigation of brain-wide stroke pathophysiology,^21^ while *in vivo* investigation of synapses in stroke has mainly focused on their role in mid-to-long-term functional adaptation and rehabilitation.^22–27^ As a result of this, relevance of synaptic plasticity in early stroke remains poorly understood.

To fill this critical gap in knowledge, we endeavored to chart in detail the earliest stages of synaptic plasticity during ischemic stroke, using immunohistochemistry, proteomics, and RNAseq. In doing so, we compared 3 key brain regions relevant to the stroke context: the **ischemic core** directly affected by blood flow disruption, the surrounding **penumbra** subjected to mild ischemia, and the **contralateral** cortex. Our findings demonstrate emergence of distinct area-specific programs of synaptic remodeling within 1 hour of stroke onset, revealing a hitherto unappreciated rapid-response brain mechanism.

## Results

### Ischemia induces rapid localized decline of synapses in neuronal cultures and *in vivo*

We firstly investigated the effect of oxygen-glucose deprivation (OGD), a canonical cell-based model of ischemia,^28,29^ on synapses formed in neuronal cultures. In line with previous evidence,^28^ OGD induced significant cell death over the course of 2 hours, indicating rapid advent of ischemia^30^ (data not shown). Immunostaining showed loss of punctate labeling for an excitatory postsynaptic density (PSD) marker protein Homer within 45 minutes (**Figure 1A, 1B**). Similar decrease of punctate staining was observed for a presynaptic active zone (AZ) scaffold protein Bassoon **(Figures 1A, 1C**) and for an inhibitory PSD protein Gephyrin (**Figure 1D, 1E**). The ratio of Gephyrin to Homer was significantly decreased by OGD, indicating a stronger effect of OGD on inhibitory compared to excitatory synapses (**Figure 1F**). After 2 hours of OGD, some of the synaptic markers exhibited a decrease in the number of puncta and an increase in nearest neighbor distance between the puncta (**Figure S1A-F**), consistent with synaptic loss. Thus, ischemia induces rapid disassembly of synapses in cultured neurons.

**Figure 1.**
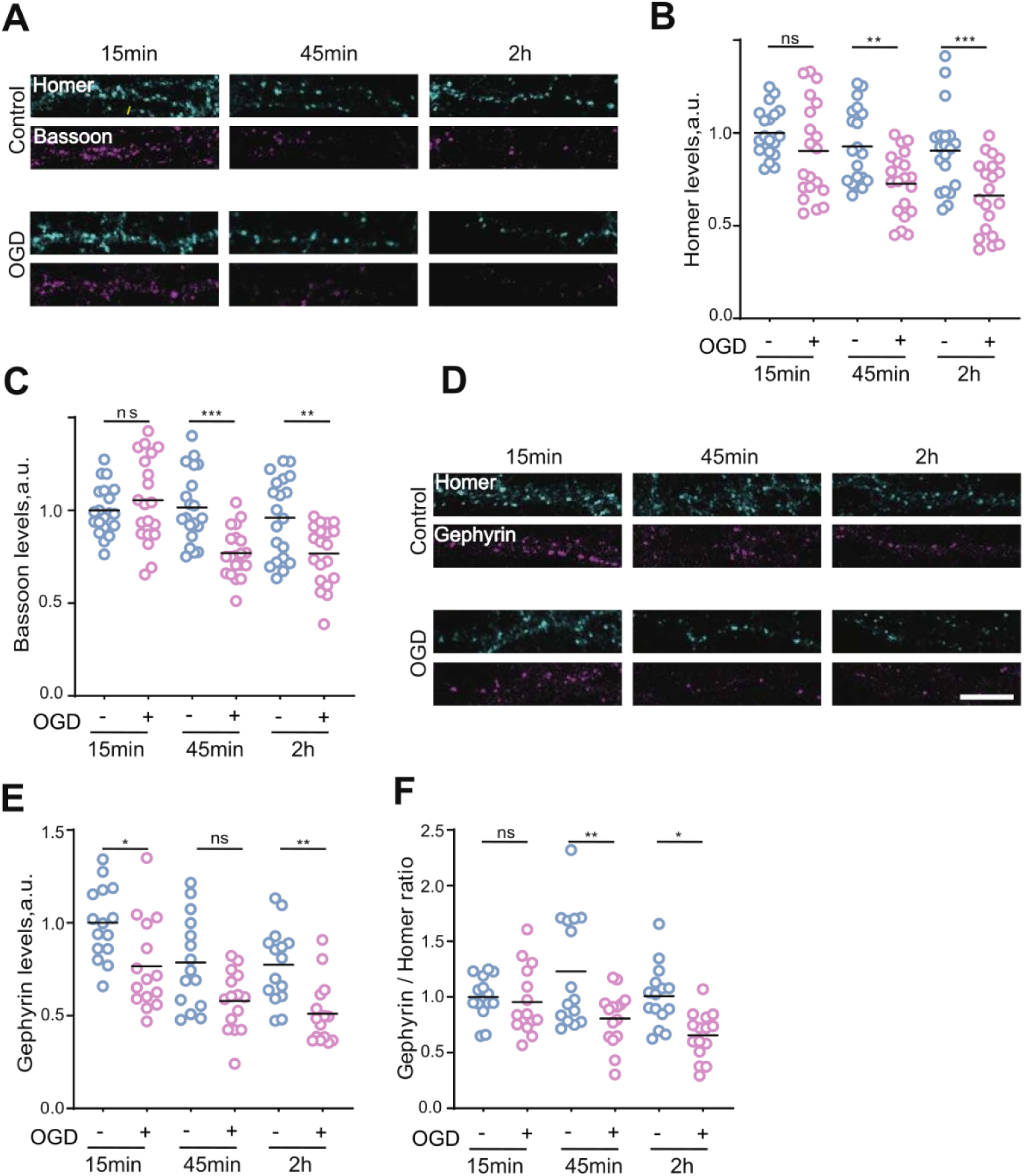
OGD induces rapid structural decline of excitatory and inhibitory synapses in cultured neurons. (A) Representative immunostaining images for Homer and Bassoon in neuronal cultures after OGD treatment. (B) Quantification of Homer in Homer(+) puncta. (C) Quantification of Bassoon in Bassoon(+) puncta. (D) Representative immunostaining images for Homer and Gephyrin in neuronal cultures after OGD treatment. (E) Quantification of Gephyrin in Gephyrin(+) puncta. (F) Quantification of Homer/Gephyrin ratio in during OGD treatment at different time points. *P < 0.05, **P < 0.01, ***P < 0.001, one-Way ANOVA analysis followed by Dunn’s multiple comparisons test and One-Way ANOVA analysis followed by Sidak’s multiple comparisons test. Scale bar, 10 μm.

Although OGD offers a convenient model mimicking the conditions inside the brain area directly subjected to ischemia, its key limitation is inability to recapitulate brain-wide of stroke physiology *in vivo*.^31^ Therefore, we leveraged the middle cerebral artery occlusion (MCAO) model in rats, which is widely considered to faithfully reproduce the morphology and physiology of human ischemic stroke, notably featuring well-defined core and penumbra areas.^31–35^ As expected, the MCAO procedure reliably produced necrotic lesions in the prefrontal cortex within 2 hours, as evidenced by neurologic examinations and Longa’s score^35^, confirming the incidence of major ischemic stroke (**Figure S2A**). In the ischemic core area, all three synaptic markers exhibited significant decline within 4 hours of MCAO (**Figure 2A-E, S2B-G**), consistent with observations in the OGD model and indicative of rapid ischemia-induced synaptic disruption of both excitatory and inhibitory synapses. Puncta of Homer and Gephyrin, but not Bassoon, were also decreased in the penumbra, suggesting some impact on PSD (**Figures 2A-E, S2B-G**). Consistent with published evidence,^36^ MCAO reduced synaptic levels of actin filaments (F-actin) in both ischemic core and the penumbra, supporting the notion of rapid synaptic remodeling (**Figure S2H-I**). Taken together, these findings provide evidence that stroke rapidly causes profound synaptic loss in the ischemic core and some synaptic decline in the penumbra.

**Figure 2.**
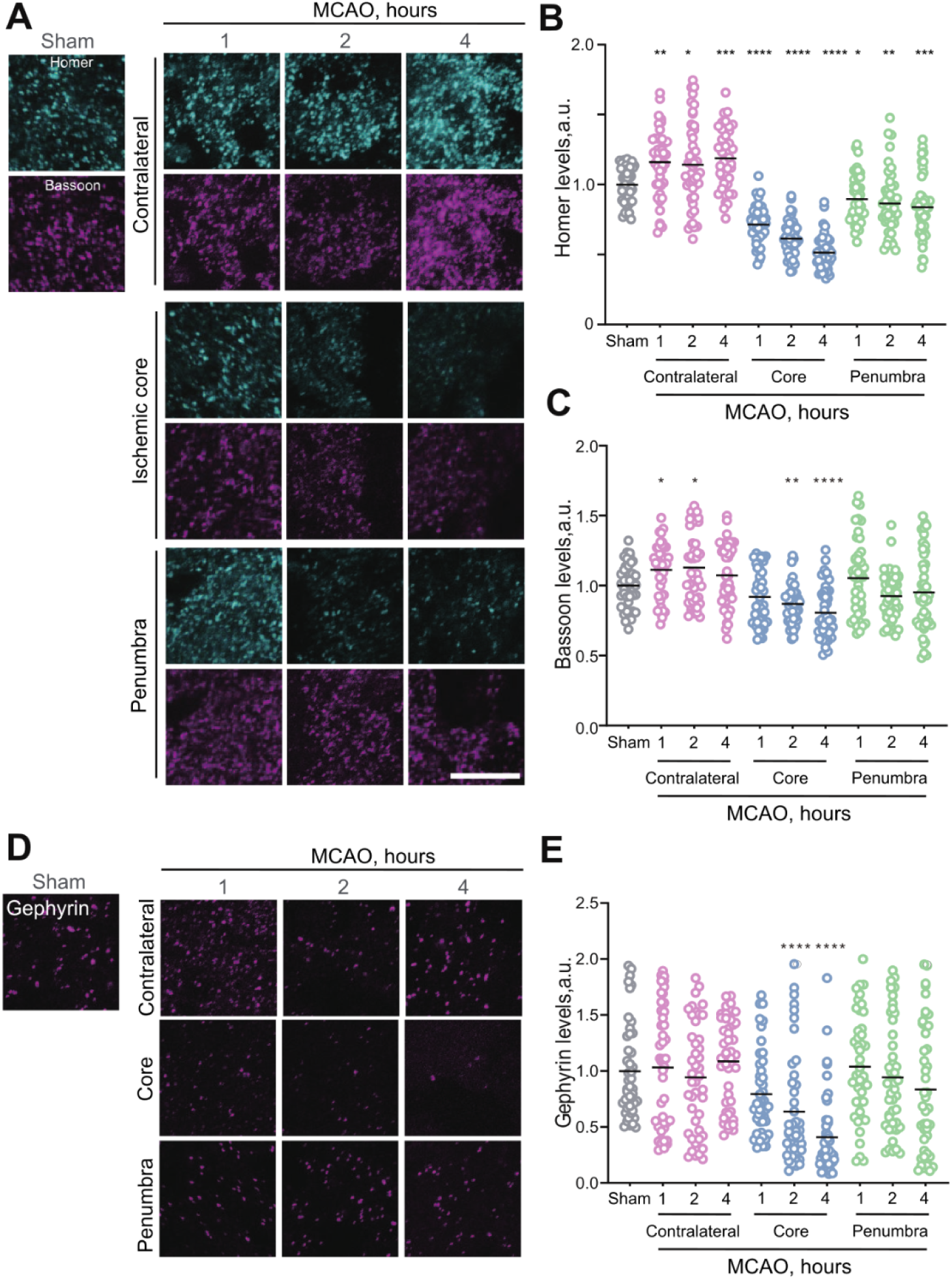
MCAO triggers rapid region-specific alteration of synaptic structure. (A) Representative immunostaining images for Homer and Bassoon immunostaining in brain sections following MCAO. (B) Quantification of Homer in Homer-positive puncta in brain sections following MCAO. (C) Quantification of Bassoon in Bassoon-positive puncta in brain sections following MCAO. (D) Representative immunostaining images for Gephyrin in brain sections following MCAO. (E) Quantification of Gephyrin in Gephyrin-positive puncta in brain sections following MCAO. *P < 0.05, **P < 0.01, ***P < 0.001, ****P < 0.0001 by One-Way ANOVA analysis followed by Dunn’s multiple comparisons test. Scale bar, 10 μm.

### Stroke triggers rapid long-range structural synaptic enhancement *in vivo*

Mid- to long-term stroke recovery is known to involve remodeling of functional connectivity across the brain.^23,27,37–39^ To investigate the long-range effects of hyperacute stroke on synaptic plasticity, we measured the effect of stroke on synaptic markers in the contralateral cortex, at a considerable distance (0.5-1 cm) from the ischemic core. Intriguingly, contralateral punctate staining for both Homer and Bassoon was significantly increased after as little as 1 hour following MCAO (**Figure 2A-C**), suggesting rapid synaptic enhancement. There was no concomitant increase in Gephyrin puncta (**Figure 2D-E**), indicating that structural synaptic enlargement was likely restricted to excitatory synapses. The numbers of Homer- and Bassoon-positive puncta were not significantly altered, consistent with MCAO-induced enlargement of existing synapses rather than generation of new ones (**Figure S2B-G**).

### Region-specific synaptic reshaping by ischemia

To get a deeper insight into stroke-induced rapid synaptic remodeling, we resorted to immunostaining for key synaptic proteins. For postsynaptic receptors, we imaged distribution of an AMPA-type glutamate receptor subunit GluA2 and a GABA receptor subunit GABRA1. OGD resulted in rapid loss of GluA2 and GABRA1 puncta within 15 and 45 min respectively (**Figure S3A-D**), consistent with previously reported effects of OGD.^40^ In the MCAO model, there was a decrease in GluA2 in both core and penumbra area, while loss of GABRA1 was only observed in the core (**Figure 3A-3D**). There were no significant changes in either GluA2 and GABRA2 in the contralateral hemisphere as detected by immunostaining (data not shown). These data indicated that ischemia and stroke induced rapid decrease in postsynaptic function.

**Figure 3.**
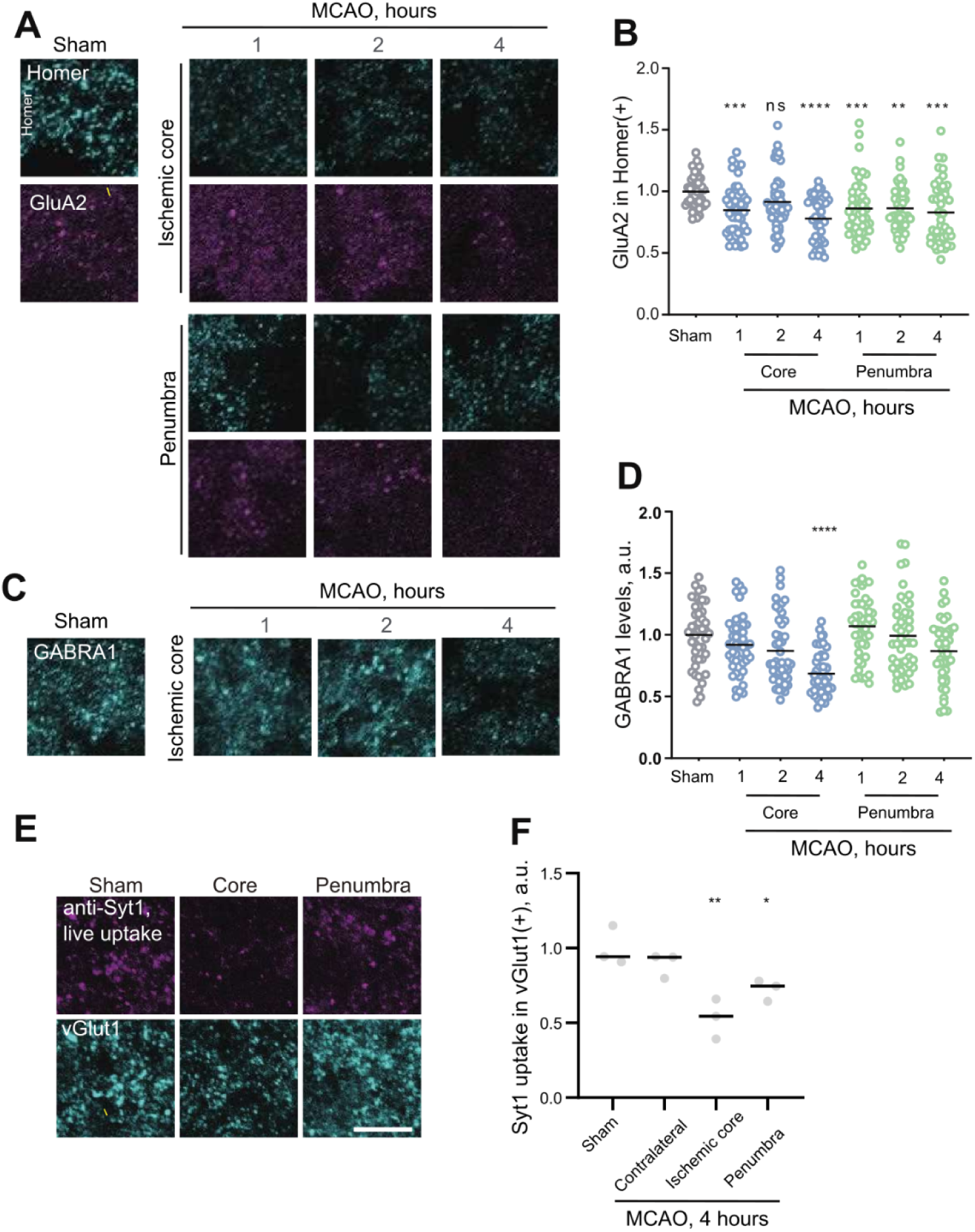
MCAO induces region-specific changes in presynaptic and postsynaptic compoments. (A) Representative immunostaining images for Homer and GluA2 in brain sections following MCAO. (B) Quantification of GluA2 in Homer(+) puncta in brain sections following MCAO. (C) Representative immunostaining images for GABRA1 in brain sections following MCAO. (D) Quantification of GABRA1 in GABRA(+) puncta in brain sections following MCAO. (E) Representative immunostaining images of acute brain slices live labeled with anti-Syt1 antibody and stained with vGlut1 post-fixation following MCAO. (F) Quantification of anti-Syt1 uptake in acute brain slices following MCAO. *P < 0.05, **P < 0.01, ***P < 0.001, ****P < 0.0001 by one-Way ANOVA analysis followed by Dunn’s multiple comparisons test. Scale bar, 10 μm.

To further investigate effects of stroke on presynaptic structure and function, brain sections were immunolabeled for P/Q-type voltage-gated Ca2+ channel (VGCCs) Ca_V_2.1, and a synaptic vesicle (SV) marker protein vesicular glutamate transporter vGlut1. We found that MCAO did not result in any change in Ca_V_2.1 staining (**Figure S3E**), while there was a significant increase in vGlut1 in the ischemic core and penumbra area (**Figure S3F**). For directly measurement of presynaptic function, we prepared acute brain sections following MCAO and performed live labeling with an antibody directed against the lumenal/extracellular domain of the SV marker protein Synaptotagmin (anti-Syt1).^41,42^ Anti-Syt1 labeling was significantly decreased in both the core and the penumbra (**Figure 3E-F**). Taken together, these findings indicated that ischemia rapidly downregulates pre- and post-synaptic function.

### Ischemia-induced NMDAR signaling regulates long-distance synaptic remodeling *in vivo*

We then sought to identify the signaling mechanism underlying long-distance synaptic strengthening. NMDA-type glutamate receptors (NMDARs) are key regulators of synaptic plasticity in disease,^43–45^ and dysregulated NMDAR signaling has been strongly implicated in neuronal death in stroke.^46^ MCAO increased staining for an essential NMDAR subunit GluN1 in both ischemic core and penumbra, but not in the contralateral area (**Figure 4A-B**). Surprisingly, this observation was at odds with the data from the OGD cell model, where no significant effect on synaptic GluN1 was observed (**Figure 4C-4D**). These findings demonstrate that although local ischemia *in vivo* rapidly recruits synaptic NMDARs, these effects cannot be recapitulated in the OGD model, highlighting the limitations of cell-based models of ischemia in recapitulating physiological complexity of stroke.

**Figure 4.**
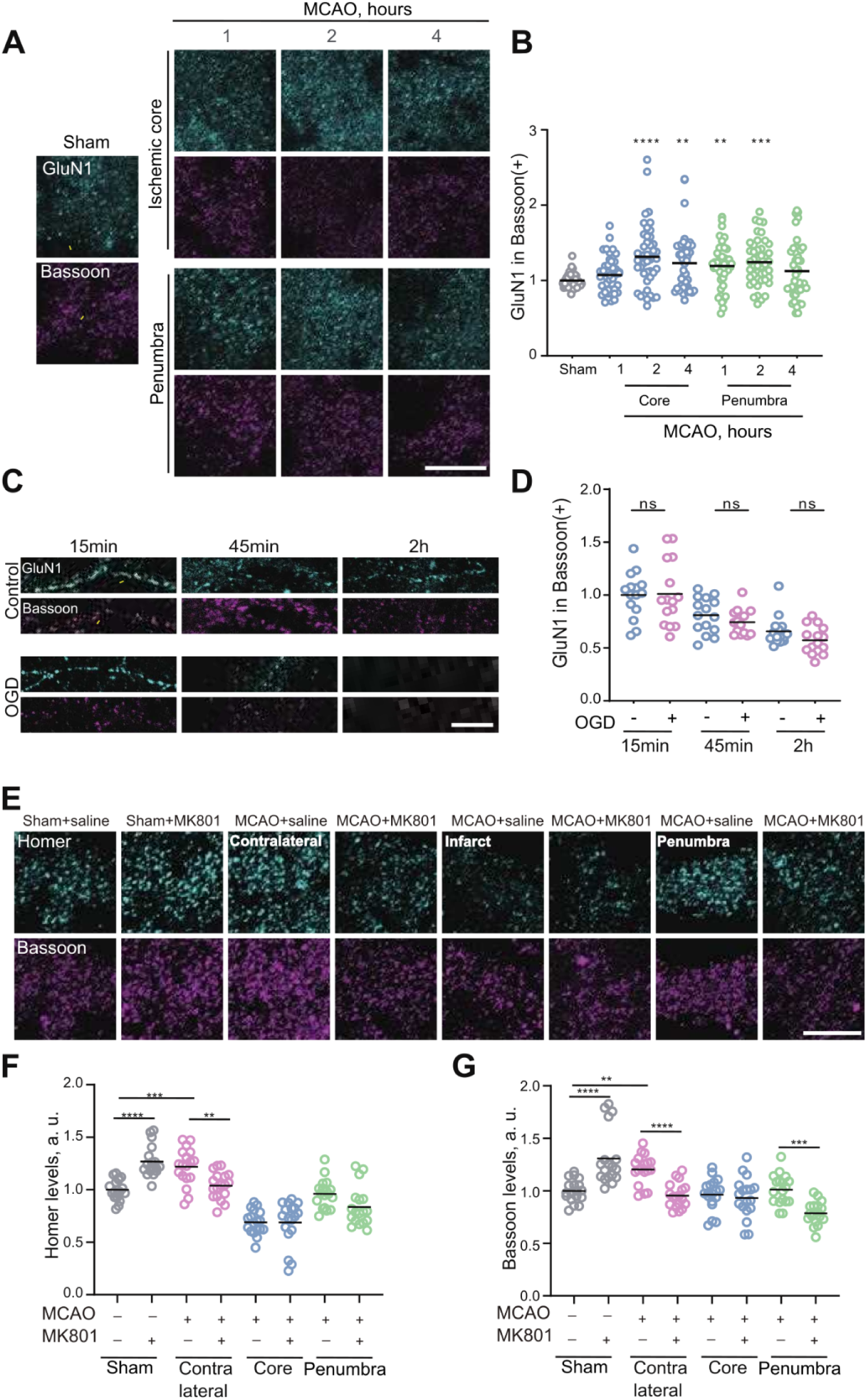
NMDAR signaling regulates rapid synaptic remodeling in the contralateral hemisphere and the penumbra during MCAO. (A) Representative immunostaining images for GluN1 and Bassoon in brain sections following MCAO. (B) Quantification of GluN1 in Bassoon(+) puncta in brain sections following MCAO. (C) Representative immunostaining images for GluN1 and Bassoon in neuronal cultures following OGD treatment on DIV14. (D) Quantification of GluN1 in Bassoon(+) puncta in neuronal cultures following OGD. (E) Representative immunostaining images for Homer and Bassoon in brain sections, taken 1 hour after MK801 injection followed by 4 hours of MCAO. (F) Quantification of Homer levels in Homer(+) puncta in brain sections, 1 hour after MK801 injection followed by 4 hours of MCAO. (G) Quantification of Bsn levels in Bassoon(+) puncta in brain sections, 1 hour after MK801 injection followed by 4 hours of MCAO. *P < 0.05, **P < 0.01, ***P < 0.001, ****P < 0.0001 by One-Way ANOVA analysis followed by Dunn’s multiple comparisons test. Scale bar, 10 μm.

To investigate the functional relevance of NMDAR signaling on synaptic enhancement in hyperacute stroke, we injected a NMDAR antagonist MK801, which has a neuroprotective effect in the MCAO model,^47^ and performed immunostaining for Homer and Bassoon following MCAO (**Figure 4E**). We found that injection of MK801 exacerbated synaptic decrease in the penumbra and abolished MCAO-induced synaptic strengthening on the contralateral hemisphere; interestingly, MK801 also increased the intensity of both Bassoon and Homer puncta in sham controls, suggesting brain-wide rapid homeostatic synaptic upscaling triggered by NMDAR blockade (**Figure 4E-G**). This evidence confirms that NMDAR signaling regulates rapid region-specific synaptic remodeling in hyperacute stroke.

### Proteomic analysis of synaptosomes in hyperacute stroke reveals systemic contralateral enhancement

To gain a broader insight into synaptic remodeling in hyperacute stroke, we leveraged quantitative proteomics to identify differentially expressed proteins in biochemically purified brain synaptosomes following 4 hours of MCAO (**Figure 5, Table S1**). Principal component analysis (PCA) showed separation between ischemic core and penumbra on one side and control and contralateral cortex on the other, suggesting that stroke effected a wide-range alteration of synaptic composition in regions directly affected by ischemia (**Figure 5A**), notwithstanding substantial variability between samples (**Figure 5B**). Synapses from the core region and penumbra exhibited considerably more downregulated *vs* upregulated proteins (580 *vs* 210 and 373 *vs* 146 respectively), indicative of general synaptic downregulation, while in contralateral synapses upregulated and downregulated proteins were almost evenly matched (88 *vs* 93), in line with the notion of synaptic remodeling. There was a degree of overlap between regional expression profiles, consistent with emergence of brain-wide as well as localized responses (**Figure 5C**).

**Figure 5.**
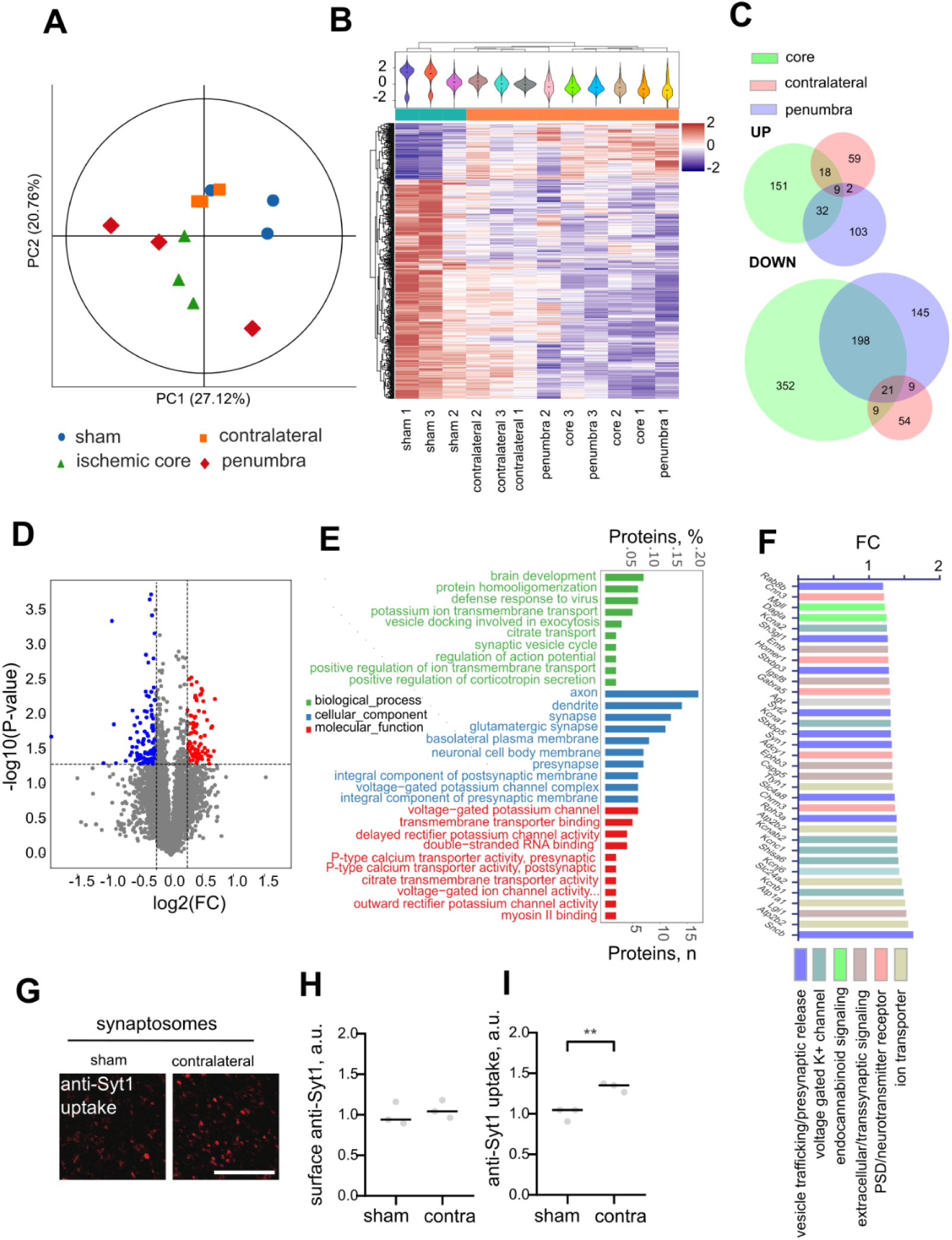
Proteomic analysis of purified synaptosomes confirms cross-brain presynaptic enhancement following MCAO. (A) PCA of protein levels for all samples. (B) Heatmap and clustering analysis for all samples. (C) Venn diagram showing numbers and overlap in upregulated (top) and downregulated (bottom) proteins between regions. (D) Volcano plot for differentially expressed proteins, contralateral hemisphere *vs* sham control. (E) Top GO terms for upregulated proteins, contralateral hemisphere *vs* sham control (F) Enrichment (FC, fold change) of synaptically relevant proteins upregulated in the control hemisphere *vs* sham control; colors highlight different functional attributions. (G) Representative immunostaining images showing KCL-induced uptake of anti-Syt1 antibody in synaptosomes from the contralateral hemisphere *vs* sham control. (H) Quantification of surface anti-Syt1 labeling in synaptosomes from the contralateral hemisphere *vs* sham control. (I) Quantification of KCL-induced anti-Syt1 uptake in synaptosomes from the contralateral hemisphere *vs* sham control. P<0.01, 2-tailed t test, N=3 animals/condition. Scale bar, 10 μm.

We focused on contralateral synapses in more detail (Figure 5D). Gene ontology (GO) analysis manifested enrichment in terms related to biological processes and molecular functions relevant to synaptic plasticity, while top cellular component terms were consistent with synaptic location (Figure 5E). These results were confirmed by gene set enrichment analysis (GSEA), revealing significant relevance for synaptic vesicle cycle (Figure S4A-C). In contrast, GO terms for downregulated proteins were more diverse, generally pertaining to metabolism and membrane trafficking, with notable enrichment for nucleoplasm in the cellular component category, suggesting a non-synaptic response likely located in the nucleus (Figure S4D). Among the 88 upregulated proteins, 35 had documented synaptic relevance as indicated by their GO terms (Figure 5F), including postsynaptic function, presynaptic release, ion transport, endocannabinoid signaling, and extracellular adhesion, with a particular increase in voltage-gated K^+^ channels, key regulators of neuronal excitability and synaptic plasticity.^48^ The most increased protein was β-Synuclein, previously suggested as a biomarker for stroke.^49,50^ Another noteworthy effect was a 40% increase in vimentin, the main protein of intermediate filaments, implicated in myelination and neuroprotection by glia in stroke. ^51^ Taken together, this evidence confirms broad synaptic enhancement in the contralateral hemisphere during hyperacute stroke.

Conversely, no synaptic enhancement was observed in the ischemic core and penumbra (**Table S1**, **Figure S5**). In core samples, upregulated GO terms were enriched for plasma membrane localization, while downregulated terms were nucleus and cytoplasm, possibly indicating contamination by disintegrated cell compartments due to ischemia-induced necrosis (**Figure S5C-D**). In the penumbra synapses, GO analysis indicated upregulation in beta-oxidation of fatty acids in mitochondria and translational downregulation in the cytosol (**Figure S5E-F**). This effect is consistent with rapid advent of a metabolic response involving reduced protein biosynthesis and increased lipid metabolism, previously demonstrated in ischemia.^52–55^

To directly confirm presynaptic enhancement in the contralateral cortex, we applied the live labeling anti-Syt1 assay to synaptosomes. Given our previous evidence showing no increase in constitutive SV cycling (**Figure 3E-F**), we resorted to depolarization by high K+ to measure evoked SV cycling.^56^ Surface labeling of synaptosomes with anti-Syt1 showed no differences between sham control and MCAO, indicating that surface levels of Syt1 were unchanged, however depolarization by 30 mM KCl resulted in a significantly higher live labeling signal in synaptosomes purified from the contralateral MCAO cortex, indicating an increase in evoked SV cycling (**Figure 5G-5I**). These data confirm that stroke triggers rapid long-distance enhancement of presynaptic function across the brain.

### Region-specific gene expression programs in synapses during hyperacute stroke

Our data revealed a pattern of region-specific synaptic reorganization in response to hyperacute stroke. To reveal associated gene expression dynamics, we isolated, sequenced and analyzed total RNA from the whole brain tissue and from synaptosomes. PCA of the data from whole brain tissue exhibited substantial scatter, consistent with inter-individual variability of brain gene expression^57^ (**Figure S6A**). Conversely, PCA of synaptosome samples returned clear separation between control, ischemic core and penumbra, suggesting emergence of large-scale, distinct, region-specific gene expression programs at the synapse triggered by stroke, while control and contralateral samples overlapped, suggesting little difference in gene expression profiles (**Figure 6A**).

**Figure 6.**
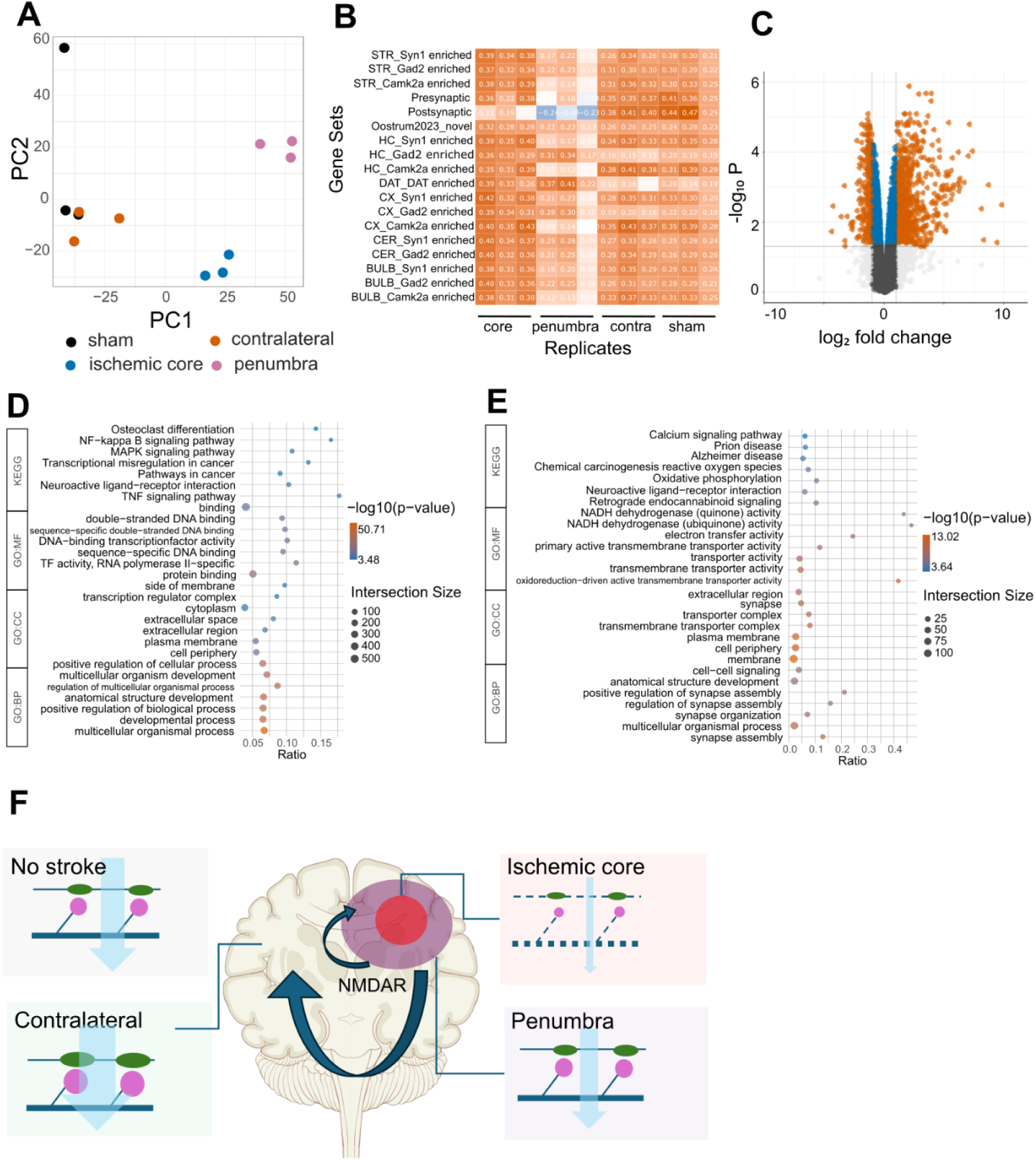
RNAseq reveals rapid induction of region-specific synaptic gene expression programs following MCAO. (A) PCA of gene expression for synaptosome samples. CA – control animals, 4z – 4 hour contralateral, 4p – 4 hour penumbra, 4c – 4 hour (ischemic) core. (B) GSVA for synaptic gene sets in synaptosomal fractions. (C) Volcano plot for differentially expressed genes in synaptosomes, penumbra *vs* sham control. (D) Functional enrichment analysis for top upregulated genes in synaptosomes, penumbra *vs* sham control. (E) Functional enrichment analysis for top downregulated genes in synaptosomes, penumbra *vs* sham control. (F) Schematic model for region-specific synaptic remodeling during hyperacute stroke. In the **ischemic core**, severe energy deficit leads to cell death and synaptic collapse. In the surrounding **penumbra,** mild ischemia triggers a response to preserve synaptic structure in an NMDAR-dependent manner, while energy-intensive synaptic function is scaled down. Across the brain, in the **contralateral hemisphere**, a long-distance relay mechanism dependent on NMDAR signaling induces homeostatic synaptic upscaling to compensate for loss of functionality in ischemic core and penumbra.

We first performed gene set variation analysis (GSVA) in synaptosomes, using previously published synaptic gene sets^58^ (**Figure 6B**). No changes were observed in samples from the contralateral hemisphere, suggesting that synaptic enhancement in the contralateral hemisphere was likely not driven by local gene expression. Conversely, samples from the ischemic core manifested specific downregulation of postsynaptic sets, while the majority of synaptic sets were downregulated in synaptosomes from penumbra, indicating an effect on local synaptic gene expression. The effect on synaptic gene sets was also evident in the whole-brain samples from penumbra (**Figure S6B**).

For further insight into biological processes regulated by gene expression in early stroke, we used GO analysis using the terms from Gene Ontology (GO:BP, GO:MF and GO:CC) and KEGG. In the penumbra, upregulated genes were involved in regulation of various biological processes, mainly related to development and immune response, highlighting broad reorganization in response to stroke, while downregulated terms included synapse and cell-cell communication, consistent with GSVA results (**Figure 6C-E, Figure S6C-S6E**). In the contralateral hemisphere, synaptosomes and whole brain tissue featured upregulation of genes pertaining to metabolic processes and complement system respectively (**Figure S7**). In the ischemic core, downregulated genes from whole brain tissues were enriched for transcription-related terms, while downregulated genes in synaptosomes were enriched in terms pertaining to synapses and cell adhesion (**Figure S8**). Taken together, our data reveal a wide-ranging programme of region-specific gene expression triggered by hyperacute stroke.

## Discussion

Investigation of synaptic plasticity in stroke has mainly focused on the recovery stage, spanning weeks to months and years.^4,23,24,27^ Here, we use a variety of methods to show that wide-ranging synaptic remodeling across the brain begins much sooner - indeed, as soon as within 1 hour of stroke onset. Thus, our results demonstrate the ability of the brain to rapidly implement broad systemic changes to functional circuitry in response to acute metabolic perturbation.

We propose that simultaneous curbing of synaptic function in the penumbra and enhancement in the contralateral cortex represents an early-stage response mechanism to safeguard brain-wide functionality in face of localized synaptic collapse at the ischemic core (**Figure 6F**). Widespread synaptic enhancement across the cortex suggests that this form of plasticity is likely of homeostatic nature, with the timescale similar to that of ketamine- and lithium-induced plasticity.^59^ It remains to be seen how rapid synaptic remodeling fits alongside other long-range mechanisms of neuronal plasticity in stroke, including excitability, functional connectivity, and neurogenesis.^38,60,61^

Our proteomic and transcriptomic analyses highlight specific functionalities associated with long-range synaptic enhancement, including membrane potential, retrograde signaling and presynaptic function, the latter directly confirmed experimentally (**Figure 5G-I**). Considering the limited timescale of our investigation, the 88 upregulated proteins identified in our survey may be mere harbingers of subsequent broader changes. Given minimal gene expression changes observed in the contralateral hemisphere, synaptic enhancement is likely triggered in a cross-brain manner from the ischemic core and the penumbra, where we find a transcriptional and translational response centered around metabolic reorganization and synaptic downregulation (**Figures 6B-E, S5**). Of note, rapid shift towards fatty acid oxidation parallels evidence from the heart, suggesting broad commonalities in metabolic adaptations to ischemia.^55^ Given the substantial energy demands of synaptic transmission,^62^ it is tempting to speculate that regional synaptic remodeling and metabolic switches may act in concert to balance functionality with tissue viability in stroke (**Figure 6F**).

Our evidence shows that NMDAR signaling orchestrates brain-wide compensatory synaptic response in early stroke (**Figure 4**), revealing a new important facet of the complex role played by NMDARs in brain disease.^43^ Rapid synaptic recruitment of NMDARs in ischemic core and penumbra is consistent with induction of NMDAR expression by global cerebral ischemia,^63^ although notably this finding could not be reproduced in the OGD system, highlighting intrinsic limitations of modeling *in vivo* stroke pathology in cell culture.^21,31^ Synaptic NMDAR recruitment is likely to be linked to ischemia-induced aberrations in neuronal activity, with possible triggers being postsynaptic depolarization^64^ or presynaptic silencing.^65^ Our evidence however does not allow to specifically assign this function to synaptic rather than extrasynaptic^44^ or indeed non-neuronal NMDARs,^45^ and the exact mechanism of the long-distance relay therefore remains unclear for now, warranting further investigation.

Owing to the limitations of our study, important questions arise concerning synaptic plasticity in early-stage neurological disorders. How is metabolic response coupled with synaptic remodeling? What is the general role of synaptic plasticity in cerebrovascular pathology, *e.g.* transient ischemic attack, traumatic brain injury, or haemorrhagic stroke? Are there any long-term functional consequences of synaptic remodeling? How is stroke-induced synaptic dynamics affected by aging?^66^ While direct investigation of synaptic remodeling in human stroke is likely to be experimentally challenging,^67^ our findings provide key novel insights into the synaptic physiology of stroke’s “golden hour” period,^5–7^ with significant clinical implications. It can be hoped that in the longer term, targeting synaptic plasticity may provide a much-needed new therapeutic modality in hyperacute stroke, paralleling emerging synaptic therapies for early treatment of neurodegenerative and neurodevelopmental disorders.^13,14,68,69^

## Acknowledgments

OG is funded by the Lewy Body Society (OOG2019/2020) and the National Natural Science Foundation of China (32070772).

Figure 6F is based on a free image titled “brain coronal section” (https://commons.wikimedia.org/wiki/File:Brain_human_coronal_section.svg) by Patrick J. Lynch, medical illustrator, and C. Carl Jaffe, MD, cardiologist.

## Author contributions

HC and YW equally contributed to this work.

Conceptualization, OG; Methodology, OG, LR, MS, QW; Investigation, HC, WY; LR; Formal Analysis, HC, WY, LR, MS, OG; Writing – Original Draft, OG, LR, MS, HC; Writing – Review & Editing, OG, MS, LR, HC, QW; Data Curation, HC, OG, LR, MS; Resources, OG, MS, QW; Supervision, OG, MS, QW; Project administration, OG, MS, QW; Funding Acquisition, OG, MS, QW.

## Methods

### Experimental models

#### Cell culture

Cultures of primary cortical neurons were derived from embryos at 16-18 days of gestation, which were isolated from pregnant rats anesthetized with isoflurane. Cerebral cortices were isolated, washed and centrifuged at 900 rpm at room temperature. Brian tissue was digested with trypsin for 20 minutes in 37°C, and digestion was terminated by the addition of DMEM containing serum, filtered through a strainer and broken down into cells by careful pipetting. Cell suspension was pelleted by gentle centrifugation, washed and resuspended in culture medium (Neurobasal medium, 2% B27 supplement, 1% glutamine). Cells were plated into 24-well plates at a density of approximately 40 000 cells per well. Medium was exchanged within 24 hours of plating, and half-changed every three days thereafter.

#### Animals

The adult male Sprague-Dawley rats (weight 230g-250g) were purchased from Jinan Peng Yue Laboratory Animal Breeding Co. Ltd and were kept under a 12-hour light/dark cycle in the same colony room with appropriate temperature. All animal experiments were conducted in compliance with National Institutes of Health Guidelines and were approved by the institutional animal care and use committee of Qingdao University.

### Method details

#### OGD

OGD was induced in 14 DIV cortical neuronal cultures with a HEPES-buffered solution (25 mM HEPES, pH 7.4, 140 mM NaCl, 5 mM KCl, 2 mM CaCl_2_, 1 mM MgCl_2_, and 10 mM sucrose or supplemented with 10mM glucose for control). Neuron culture plates were washed twice and incubated with OGD-HEPES solution in an anaerobic workstation (Ruskinn Concept 400, United Kingdom.) at 37°C, 95% N_2_ and 5% CO_2_ prior to being harvested or fixed at specific time points (15minutes, 45 minutes and 1 hour.). Control neurons were incubated at 37°C, 5% CO_2_ with control HEPES solution and harvested or fixed at the same time points as those subjected to OGD.

#### MCAO

The skin was cut with scissors after disinfection in the middle of the neck, and the right common carotid artery (CCA), external carotid artery (ECA), and internal carotid artery (ICA) were isolated with forceps. And all were ligated with surgical sutures, a small incision was cut at the distal end of the ECA, wherein a filament was inserted. The sutures of the ICA were then undone and the inserted filament was pushed from the right CCA into the right ICA, resulting in occlusion of the origin of the right middle cerebral artery (MCA). Following the procedure, animals were sacrificed after 1, 2, and 4 hours. For sham control, the procedure was followed as described above, except no filament was inserted.

#### Proteomics

Animals were divided into two groups, MCAO and sham control, with 3 animals in each group. From MCAO animals, brain tissue was excised from each of the following regions: the ischemic core, the penumbra and the contralateral hemisphere. From sham control animals, brain tissue was excised from the cortex. The total number of samples was therefore 12, with 3 samples each in: sham control, ischemic core, penumbra and contralateral hemisphere.

300 mg of the brain tissue from each sample was homogenized in 10 volumes of the Syn-PER Reagent following manufacturer’s instructions, with 10 up-and-down even strokes using a Dounce tissue homogenizer. Homogenized tissue was centrifuged at 1200 × g for 10 minutes to remove the debris, and the supernatant was centrifuged at 15,000 × g for 20 minutes. The pellets, containing synaptosomes, were gently resuspended in the reagent.

Tandem Mass Tagging (TMT) proteomic analysis of the synaptosomal fraction was performed and analyzed by Oe Biotech (Shanghai, China) as briefly described below. One part of the synaptosomes was used for protein concentration determination and SDS-PAGE detection, and the other part was trypsinized and labeled. An equal amount of each labeled sample was then mixed and separated by chromatography. Samples were analyzed by LC-MS/MS and data analysis. Proteins were considered differentially expressed when the following conditions were met: p value <0.05, fold change (FC) <0.8 and >1.2. Overlap in protein expression profiles between different regions was visualized using DeepVenn.^70^ GSEA for proteins was carried out using WEB-based GEne SeT AnaLysis Toolkit^71^.

#### Live anti-Syt1 uptake assay

For anti-Syt1 uptake in acute brain sections, 3 animals were subjected to MCAO, while 3 animals were used as sham controls. Animals were sacrificed 4 hours after surgery. Sections of 200 μm in thickness were prepared using vibratome and allowed to recover for 30 mins following sectioning. Sections were live-labeled with anti-Syt1 antibody (1/200) for 30 mins at 37°C. Section were then washed, fixed for 20 mins in 4% PFA, permeabilized and blocked for 1.5 hours, incubated with anti-vGlut1 primary antibody overnight at 4°C. Next day, slices were rewarmed to RT for 75 mins, washed for 5 mins 6 times, then incubated in the mix of secondary antibodies for 2 hours at RT. Slices were then washed for 5 mins 6 times and mounted onto slides to be stored at 4°C.

For anti-Syt1 uptake in synaptosomes, the latter were isolated as described above. Aliquoted synaptosomes were resuspended and live labeled with 1/200 anti-Syt1 at 4 °C for 20 minutes. Synaptosomes were either fixed immediately to measure surface Syt1 or incubated with 30 mM KCL for 15 min at 37° to measure depolarization-evoked SV cycling. Synaptosomes were pelleted by centrifuging for 5 minutes at 3700 rpm and washed twice with PBS, then fixed with 4% paraformaldehyde (PFA) in PBS, followed by centrifugation for 20 minutes at 3700 rpm. Fixed synaptosomes were washed for 5 minutes 4 times, permeabilized for 30 minutes, then labeled with secondary antibodies for 1 hour, washed as above, and mounted in DAPI-containing medium onto microscopy slides.

#### RNA sequencing and analysis

Total RNA was extracted by TRIzol Reagent. RNA sequencing and quality control were provided by Beijing Novogene (Beijing, China). Transcript expression from RNA-seq was quantified using Salmon^72^ and the *Rattus norvegicus* index, followed by differential analysis using a linear mixed model, with the aid of variancePartition^73^ and the Dream^74^ method. Using contrasts, we used pairwise differential gene expression analysis between conditions to identify significant upregulated and downregulated genes (*p-value* < 0.05 and fold change > 2). Upregulated and downregulated genes underwent functional enrichment analysis using gProfiler2^75^ and “GO:BP”, “GO:CC”, “GO:MF”, “KEGG”, “MIRNA” as sources. Factor analysis using custom genesets^58^ was done with GSVA^76^ over the dream normalized counts. Graphical representations utilized ggplot2^77^ and enhancedVolcano.^78^

#### Immunostaining

For immunocytochemistry, after treatment cells were fixed with 4% PFA in PBS for 20 min at room temperature (RT) and washed 5 min (3X) before being blocked in 0.3% Triton-X100 in PBS supplemented with 5% goat serum for 30 min. Blocked coverslips were incubated with appropriate primary antibodies for 60 min at RT, then washed 5 min (4X) in PBS and incubated with secondary antibodies (1:500 AlexaFluor-488 and AlexaFluor-594) each for 60 min at RT. Coverslips were washed for 5 min (4X) before being mounted on microscope slides with mounting media (containing DAPI).

For immunohistochemistry, following MCAO brain tissue was excised from each of the following regions: the ischemic core, the penumbra and the contralateral hemisphere. From sham control animals, brain tissue was excised from the cortex. Animals were anesthetized with isoflurane and transcardially perfused with PBS (pH7.4). The brains were dissected out and kept in −80°C. Brain sectioning was performed in the coronal plane at −20°C using a Cryostat (Leica CM1950), at thickness of 25 μm. Brain sections were attached to positively-charged microscope slides and fixed with 4% PFA in PBS for 20 mins at RT, followed by 3X 10 mins (3X) in PBS. Subsequent sections were collected and stored at −80°C for future use. Sections were incubated in blocking buffer (5% goat serum, 0.3% Triton-X 100 in PBS) for 1 h at RT, and incubated for 36-48 h at 4°C with primary antibodies diluted in blocking buffer. Sections were subsequently washed for 10 min (3X) in PBS, followed by incubating with Alexa Fluor 488- and Alexa Fluor 594-conjugated secondary antibodies for 75 mins at RT, and finally washed for 10 mins (3X) with PBS. Then all sections were mounted with DAPI-containing mount media.

#### Microscopy imaging

Confocal imaging was carried out using a Nikon Eclipse Ti2 microscope. Serial confocal z stack images (0.5 μm step for 2 μm at 512 x 512 or 1024 x 1024 pixels, zoom 2 or zoom 1) were acquired with a 100x/1.45 Oil objective (Plan APO λ). Pinhole size was 1 Airy unit. Excitation laser wavelengths were 488 and 561 nm. Bandpass filters were set at 500−550 nm and 570-620 nm for imaging Alexa Fluor 488 and 570−620 nm Alexa Fluor 594 respectively. Other settings such as gain value were optimized to ensure appropriate dynamic range, low background and sufficient signal/noise ratio. Image acquisition settings were kept consistent within the experiment.

### Quantification and statistical analysis

#### Image Analysis

ImageJ was used for image analysis. For analysis of individual synaptic puncta, images were binarized by using the “Moments” setting, and puncta were identified automatically using the “Analyze Particles” command across the whole image. Fluorescence signal intensities were quantified for each synaptic punctum using the Region Of Interest (ROI) Manager function of ImageJ. To avoid rare overlap of multiple synapses, only ROIs with areas ranging from 0.1 to 2 μm^2^ were included in further analysis. All values of circularity were included in analysis. Individual ROIs within the image were merged into one compound ROI using the “Combine” and “Add” functions of the ROI Manager interface, whereupon quantification of mean signal intensity in each channel was performed using the “Measure” function. Since background fluorescence intensity was typically less than 1% of the median fluorescence in each channel within a ROI, background subtraction did not significantly affect the measurements and was not performed.

#### Statistics

GraphPad Prism software was used for statistical analysis. The results of the experiments were presented as scatter graphs, showing individual data points, with lines denoting mean values. All experiments were performed in at least three biological replicates as indicated in the Figure legends. Distributions were assessed for normality using d’Agostino and Pearson omnibus normality tests. For normally distributed datasets, Student’s t-test, 1-way ANOVA and Dunnett’s post-test were used to assess statistical significance as appropriate; otherwise, Kruskal-Wallis test was used.

### Key resources table

| REAGENT or RESOURCE | SOURCE | IDENTIFIER |
| --- | --- | --- |
| **Antibodies** |  |  |
| Homer 1 | Synaptic systems | 160002 |
| Bassoon | Abcam | ab82958 |
| GABA-A receptor alpha1 | Synaptic systems | 224203 |
| GluA2 | Sigma-Aldrich | AB1768-I |
| Gephyrin | Synaptic systems | 147011 |
| vGlut1 | Synaptic systems | 135302 |
| Cav2.1 | Synaptic systems | 152203 |
| Synaptotagmin1 | Synaptic systems | 105311 |
| Goat Anti-Rabbit IgG H&L / AF488 antibody | Bioss | bs-0295G-AF488 |
| Goat Anti-Mouse IgG/Alexa Fluor 594 | Solarbio | K1031G-AF594 |
| **Bacterial and virus strains** |  |  |
| **Biological samples** |  |  |
| **Chemicals, peptides, and recombinant proteins** |  |  |
| KCl | Sigma Aldrich |  |
| PFA | Sigma Aldrich |  |
| MK801 | Sigma Aldrich |  |
| Isoflurane | Sigma Aldrich |  |
| **Critical commercial assays** |  |  |
| Syn-PER assay for synaptosome purification | Fisher Scientific | 87793 |
| TRIzol assay for RNA extraction | Fisher Scientific | 15596026 |
| **Experimental models: Cell lines** |  |  |
| Primary rat cortical neurons, mixed culture | E16-E18 embryos |  |
| **Experimental models: Organisms/strains** |  |  |
| *Rattus norvegicus* | Jinan Peng Yue Laboratory Animal Breeding Co | https://www.dnb.com/business-directory/company-profiles.jinan_pengyue_experimental_animal_breeding_co_ltd.0ba8b547d8911a368fa3a9190a4d42b3.html |
| **Oligonucleotides** |  |  |
| **Recombinant DNA** |  |  |
| **Software and algorithms** |  |  |
| ImageJ v1.54g | https://imagej.net/ij/ |  |
| Inkscape 1.3.2 | https://inkscape.org/ |  |
| Salmon | https://combine-lab.github.io/salmon/about/ |  |
| DeepVenn | https://www.deepvenn.com/ |  |
| ggplot2 | https://ggplot2.tidyverse.org/ |  |
| WEB-based GEne SeT Analysis Toolkit | https://www.webgestalt.org/ |  |
| **Other** |  |  |

## Data Availability

Original data is available from the authors upon reasonable request.

## Supplemental Figures

**Figure S1.**
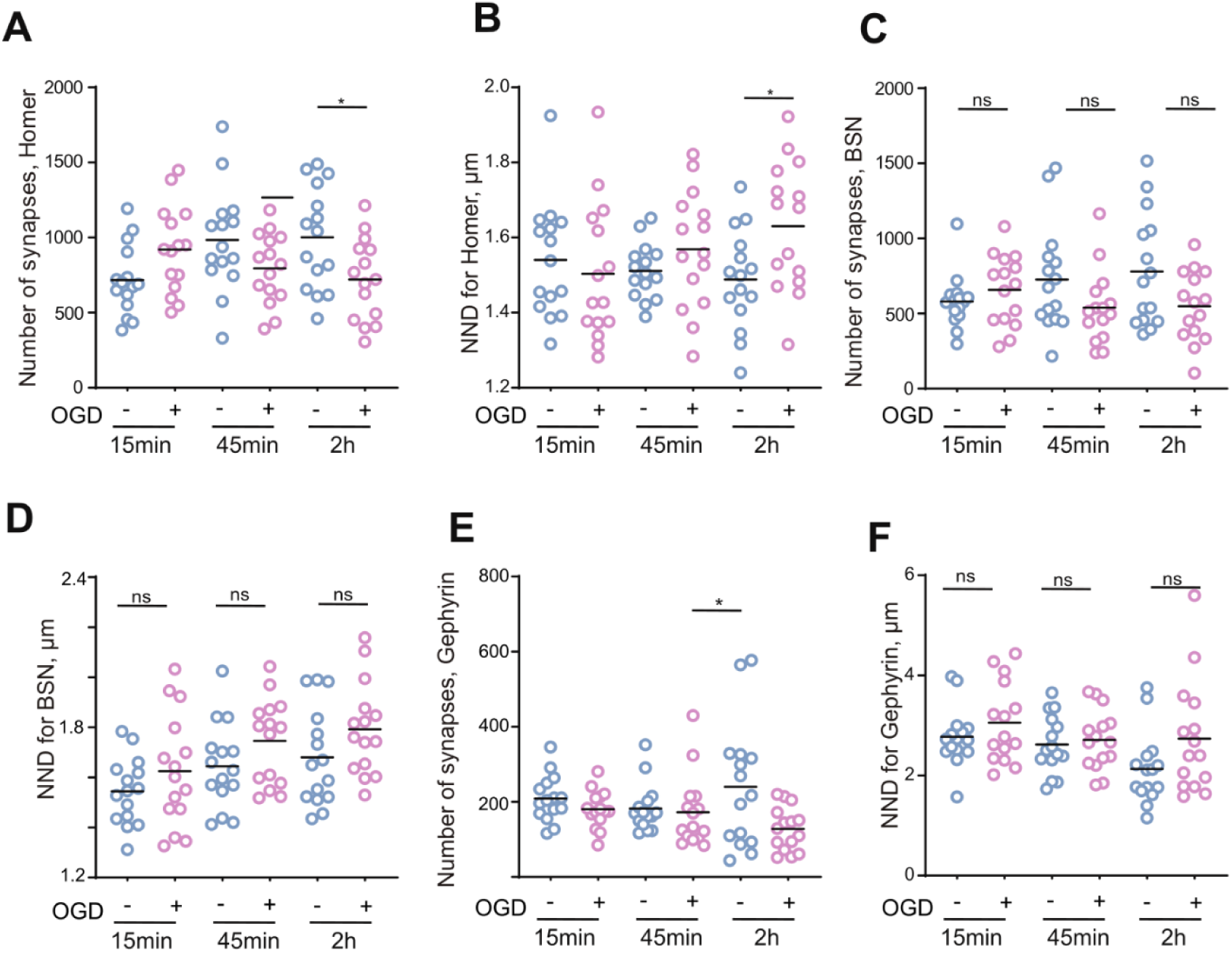
Changes in numbers and nearest neighbor distances for different synaptic markers in neuronal cultures after OGD. (A) Quantification of numbers for Homer(+) puncta. (B) Quantification of nearest neighbor distances for Homer(+) puncta. (C) Quantification of numbers for Bassoon(+) puncta. (D) Quantification of nearest neighbor distances for Bassoon(+) puncta. (E) Quantification of numbers for Gephyrin(+) puncta. (F) Quantification of nearest neighbor distances for Gephyrin(+) puncta. *P < 0.05, ns - not significant, one-way ANOVA analysis followed by Dunn’s multiple comparisons test, one-way ANOVA analysis followed by Sidak’s multiple comparisons test.

**Figure S2.**
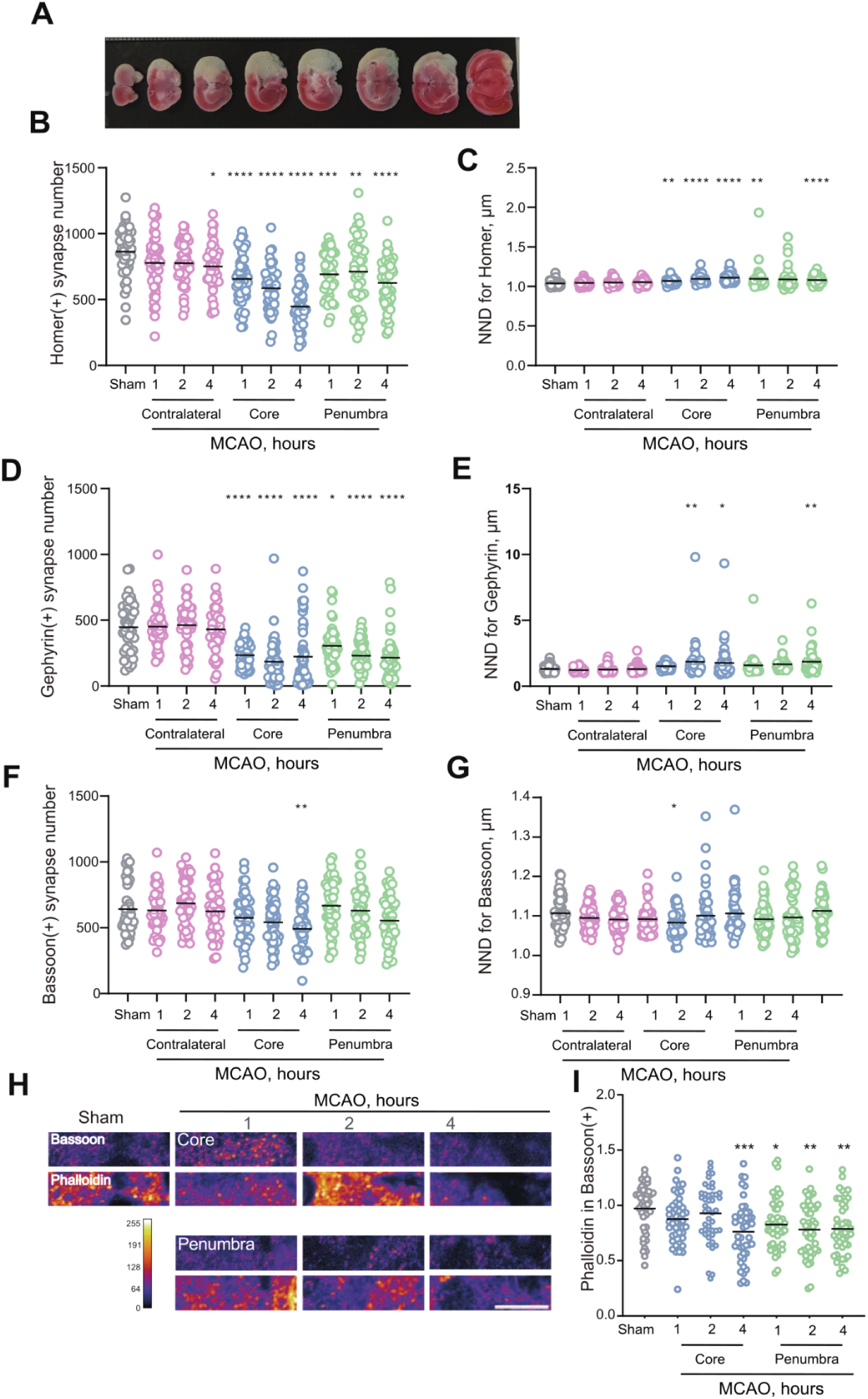
Changes in the number and nearest neighbor distance of synapses after MCAO. (A) Example TTC staining of a coronal brain section following MCAO; white part represents ischemic core. (B) Quantification of numbers for Homer(+) puncta in brain sections following MCAO. (C) Quantification of nearest neighbor distances for Homer(+) puncta in brain sections following MCAO. (D) Quantification of number for Gephyrin(+)puncta in brain sections following MCAO. (E) Quantification of nearest neighbor distances for Gephyrin(+)puncta in brain sections following MCAO. (F) Quantification of numbers for Bassoon(+) puncta in brain sections following MCAO. (G) Quantification of nearest neighbor distances for Bassoon(+) puncta in brain sections following MCAO. (H) Representative images of phalloidin and Bassoon staining in brain sections following MCAO. (I) Quantification of phalloidin staining in Bassoon(+) puncta in brain sections following MCAO. *P < 0.05, **P < 0.01, ***P < 0.001, ****P < 0.0001, one-way ANOVA analysis followed by Dunn’s multiple comparisons test. n=45 (15 animals/condition, 3 images/animal).

**Figure S3.**
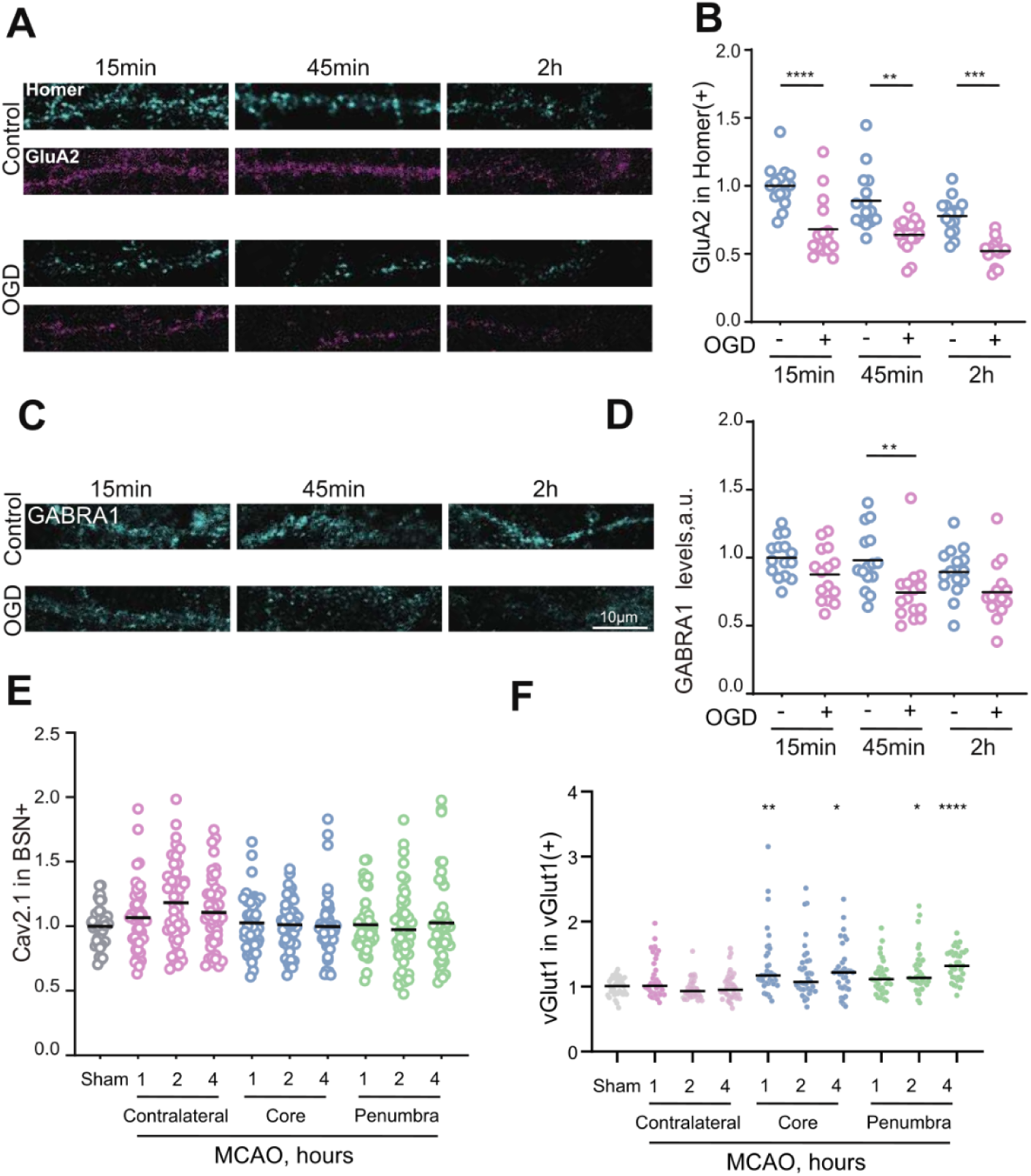
Differential recruitment of synaptic machinery during OGD and MCAO. (A) Representative immunostaining images for Homer and GluA2 in neuronal cultures after OGD treatment on DIV14. (B) Quantification of GluA2 in Homer(+) puncta in neuronal cultures following OGD. (C) Representative immunostaining images for GABRA1 in neuronal cultures after OGD treatment on DIV14. (D) Quantification of GABRA1 in GABRA1(+) puncta in neuronal cultures following OGD. (E) Quantification of Cav2.1 in Bassoon(+) puncta in brain sections following MCAO. (F) Quantification of vGlut1 in vGlut1(+) puncta in brain sections following MCAO. *P < 0.05, **P < 0.01, ***P < 0.001, ****P < 0.0001, one-way ANOVA analysis followed by Dunn’s multiple comparisons test.

**Figure S4.**
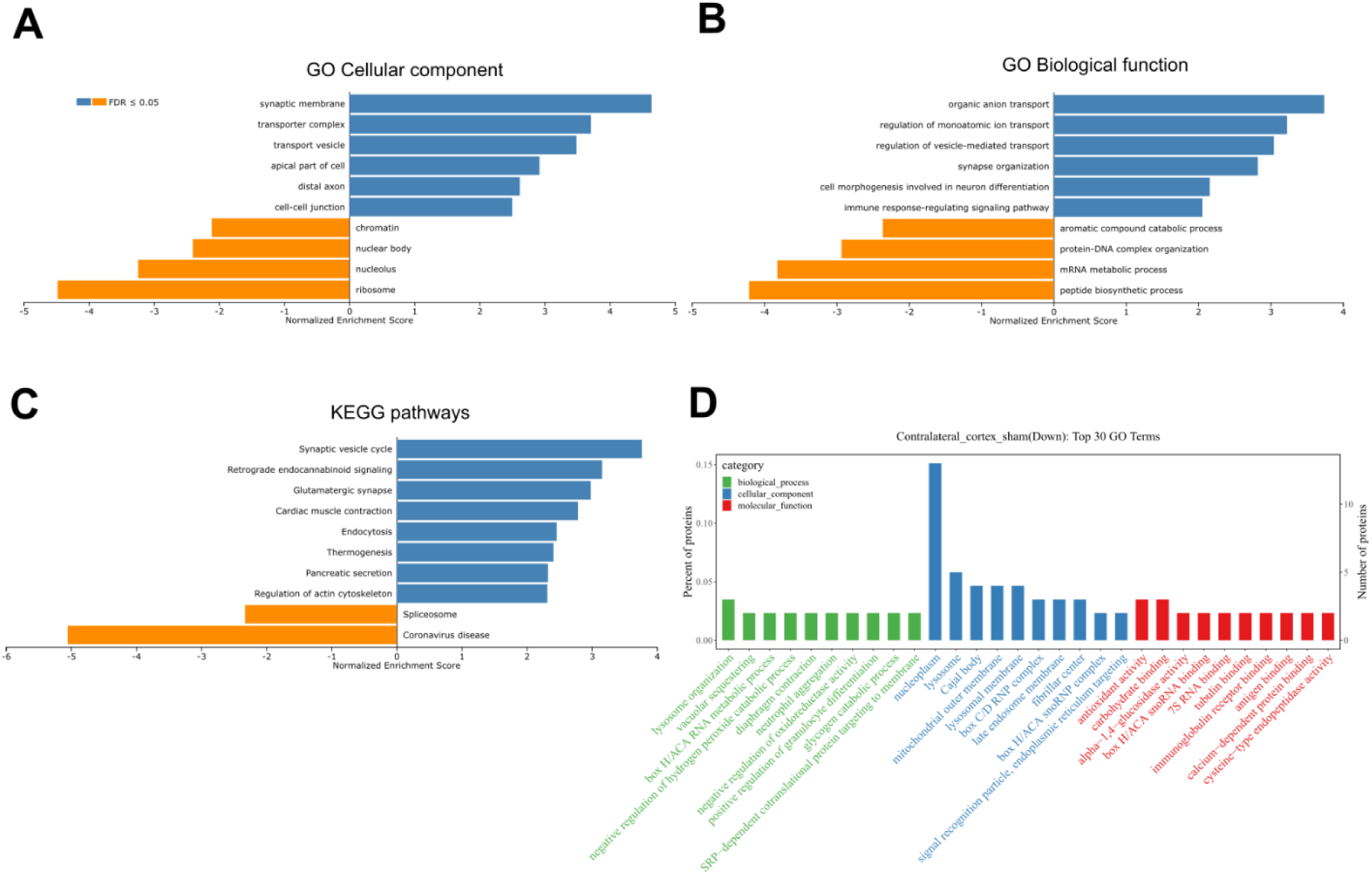
Further analysis of differentially expressed proteins in the synaptosomal fraction, contralateral hemisphere *vs* sham control. (A) Graph showing significantly enriched GO Cellular component terms. (B) Graph showing significantly enriched GO Biological function terms. (C) Graph showing significantly enriched KEGG terms for signaling pathways. (D) Graph showing top downregulated GO terms.

**Figure S5.**
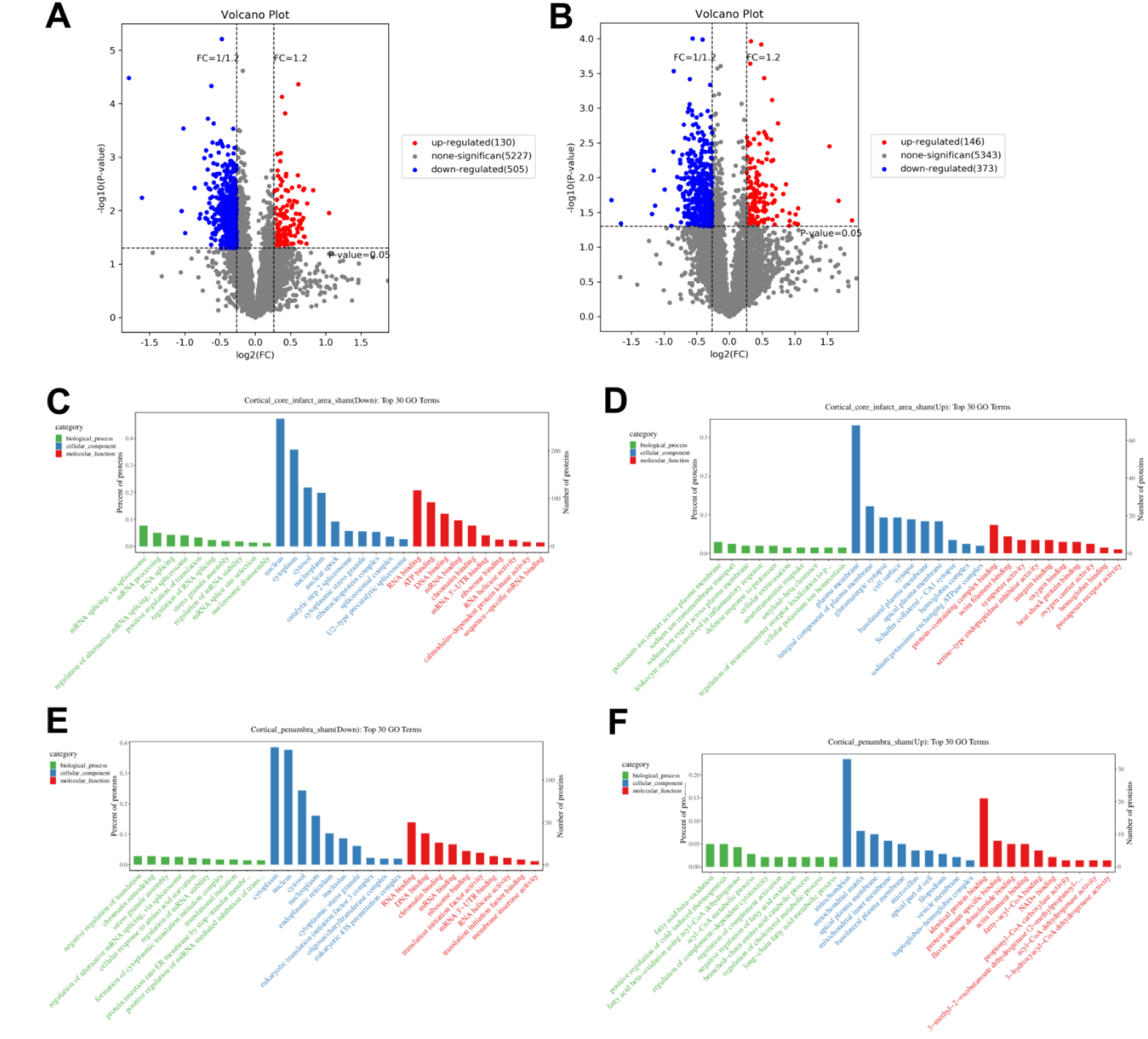
Analysis of differentially expressed proteins in the synaptosomal fraction, ischemic core and penumbra *vs* sham control. (A) Volcano plot for differentially expressed proteins, ischemic core *vs* sham control. (B) Volcano plot for differentially expressed proteins, penumbra *vs* sham control. (C) Graph showing top downregulated GO terms, ischemic core vs sham control. (D) Graph showing top upregulated GO terms, ischemic core *vs* sham control. (E) Graph showing top downregulated GO terms, penumbra *vs* sham control. (F) Graph showing top upregulated GO terms, penumbra *vs* sham control.

**Figure S6.**
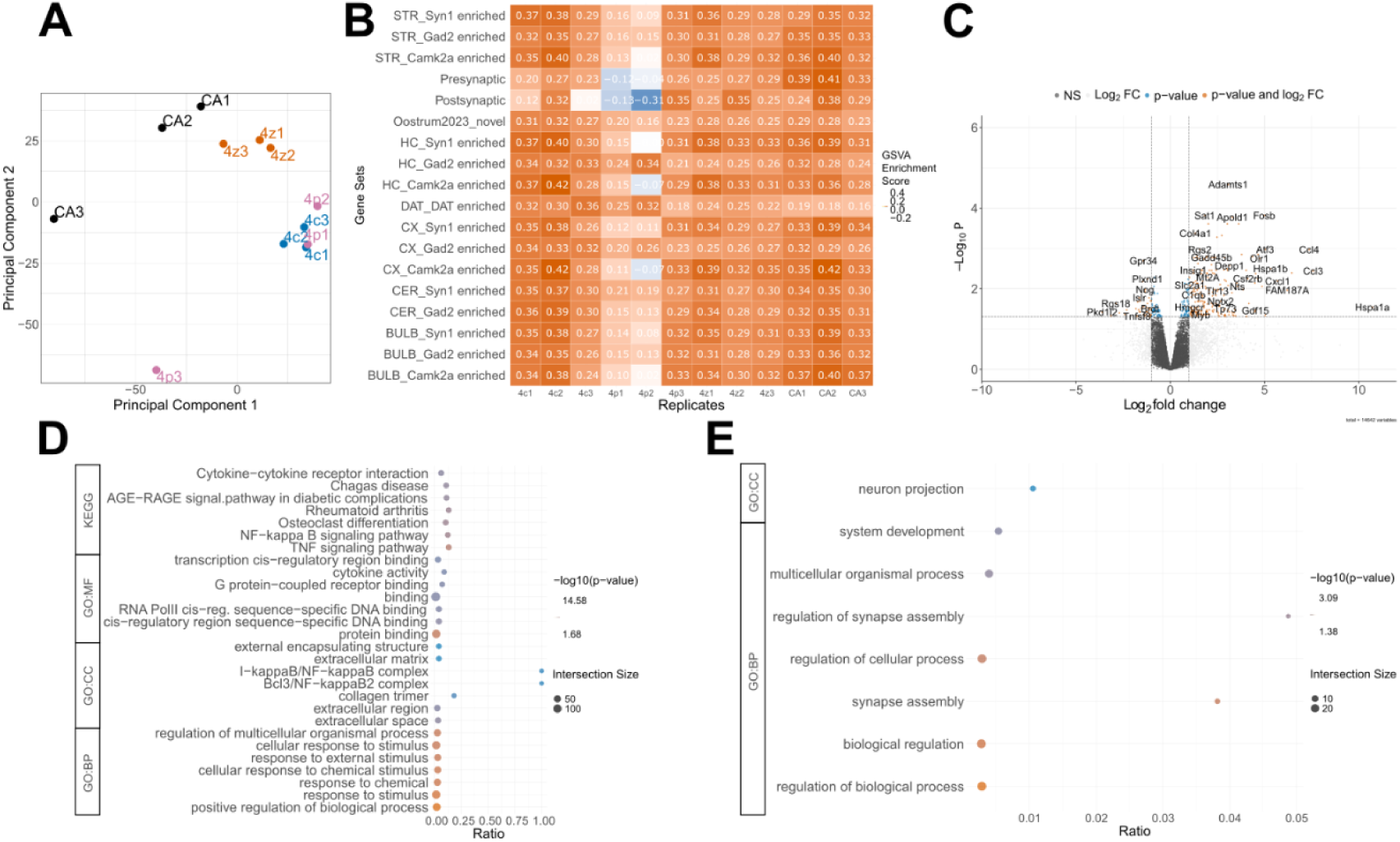
Analysis of gene expression in whole brain tissue and GO analysis of differentially expressed genes in the whole brain tissue from penumbra. (A) PCA of gene expression for the whole brain tissue. CA - sham control, z - contralateral hemisphere, c - ischemic core, p - penumbra. (B) GSVA for synaptic sets in the whole brain tissue samples. (C) Volcano plot for differentially expressed genes in the whole brain tissue, penumbra *vs* sham control. (D) Functional terms for upregulated genes in the whole brain tissue, penumbra *vs* sham control. (E) Functional terms for downregulated genes in the whole brain tissue, penumbra *vs* sham control.

**Figure S7.**
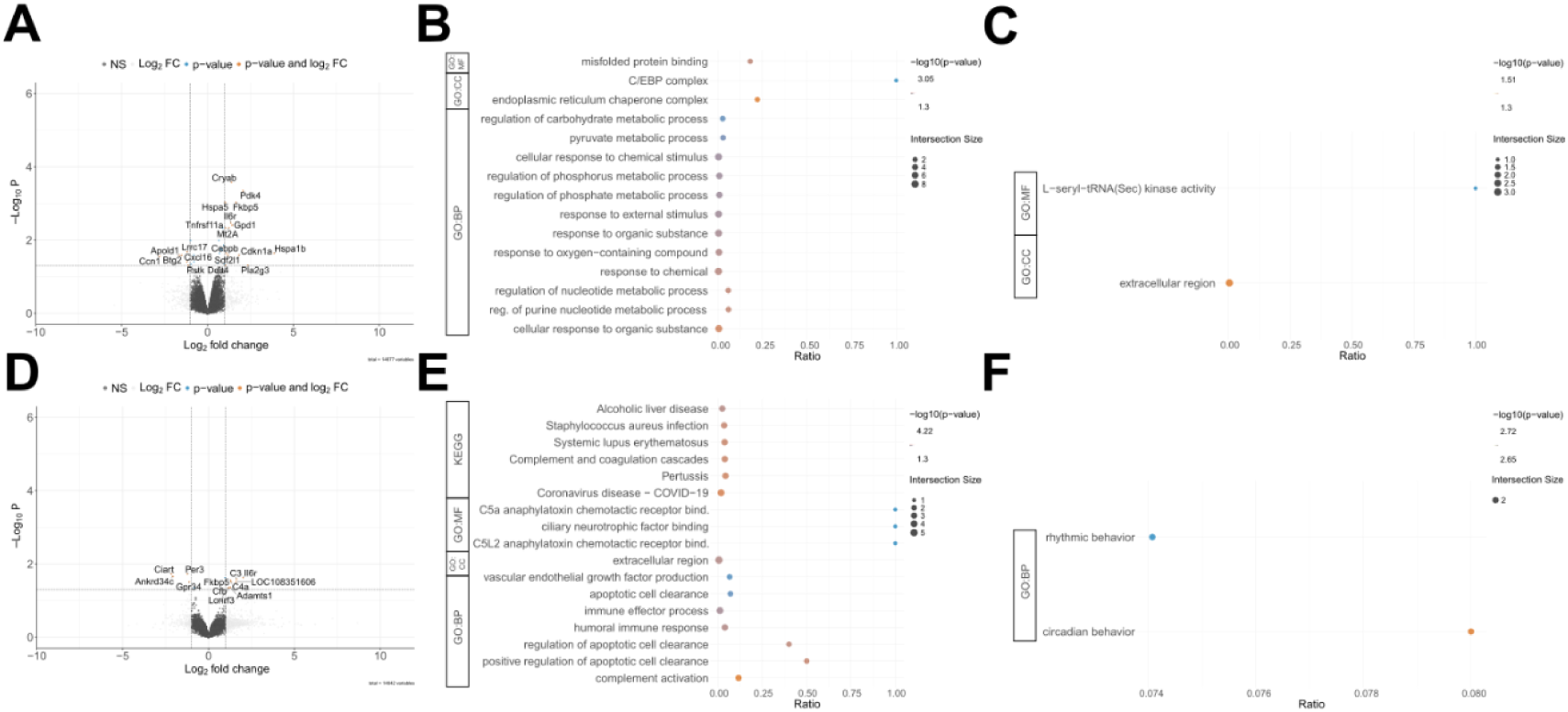
GO analysis of differentially expressed genes in the contralateral hemisphere. (A) Volcano plot for differentially expressed genes in the synaptosomal fraction, contralateral hemisphere *vs* sham control. (B) Functional terms for upregulated genes in the synaptosomal fraction, contralateral hemisphere *vs* sham control. (C) Functional terms for downregulated genes in the synaptosomal fraction, contralateral hemisphere *vs* sham control. (D) Volcano plot for differentially expressed genes in the whole brain tissue, contralateral hemisphere *vs* sham control. (E) Functional terms for upregulated genes in the whole brain tissue, contralateral hemisphere *vs* sham control. (F) Functional terms for downregulated genes in the whole brain tissue, contralateral hemisphere *vs* sham control.

**Figure S8.**
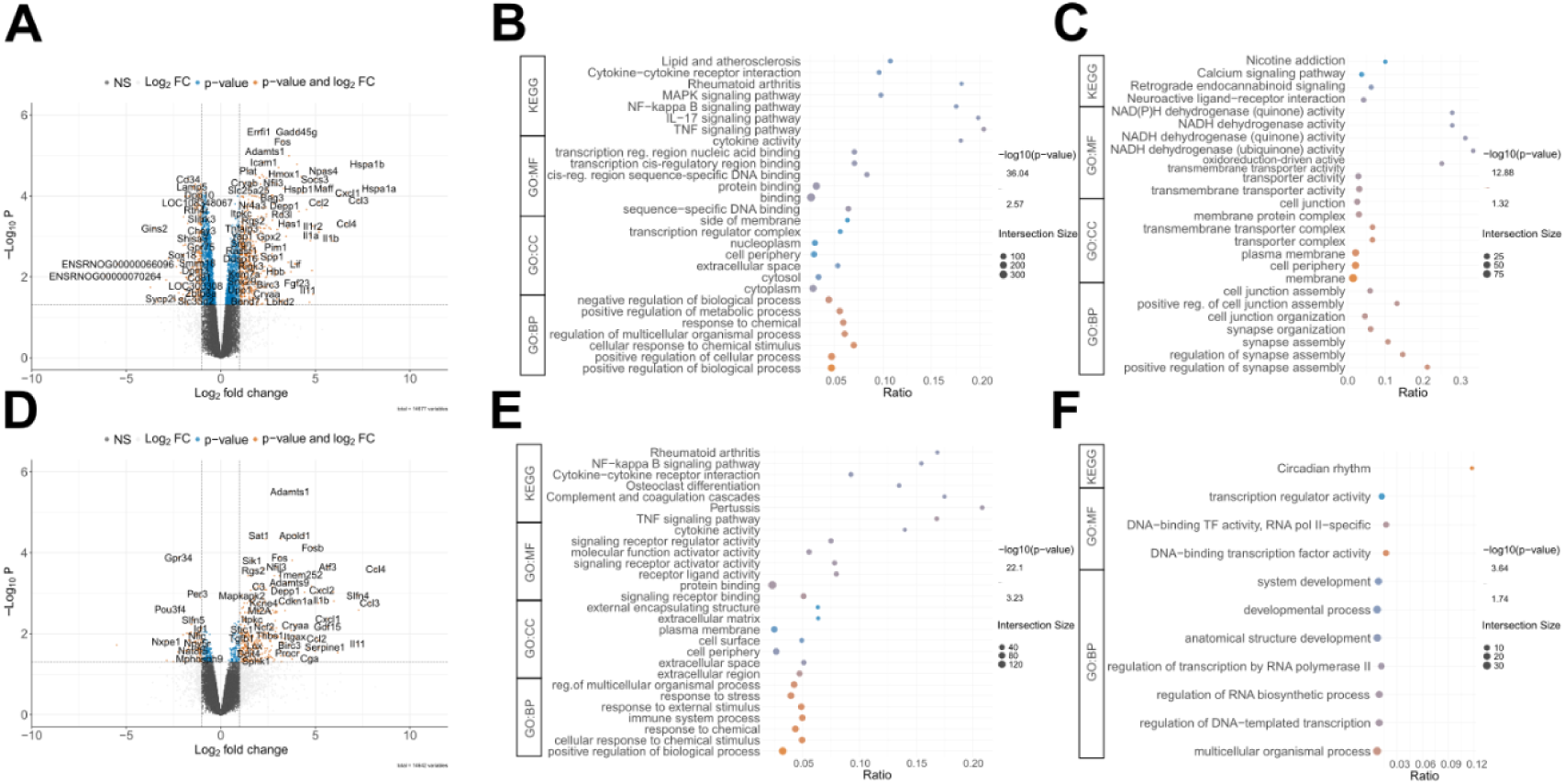
GO analysis of differentially expressed genes in the ischemic core. (A) Volcano plot for differentially expressed genes in the synaptosomal fraction, core *vs* sham control. (B) Functional terms for upregulated genes in the synaptosomal fraction, core *vs* sham control. (C) Functional terms for downregulated genes in the synaptosomal fraction, core *vs* sham control. (D) Volcano plot for differentially expressed genes in the whole brain tissue, core *vs* sham control. (E) Functional terms for upregulated genes in the whole brain tissue, ischemic core *vs* sham control. (F) Functional terms for downregulated genes in the whole brain tissue, ischemic core *vs* sham control.

## Supplemental Tables

**Table S1. Differentially expressed proteins in synaptosomes from contralateral, core, and penumbra regions, relative to the sham controls.**

**Table S2. Differentially expressed genes in whole-brain tissue and synaptosomes from contralateral, core, and penumbra regions, relative to the sham controls.**

